## Supplementary material for "Genetic manipulation of the Brassicaceae smut fungus *Thecaphora thlaspeos*": Supplementary files.pdf

Journal of Fungi

MDPI

**Table S1. Pilot study for protoplasting enzyme cocktail.** Various conditions for each enzyme combination were tested mostly in single replicates.

| Organism | Enzyme | Buffer composition | Temperature | Incubation time | Yield | Reference |
|---|---|---|---|---|---|---|
| *U. maydis* | 12,5 mg/mL Glucanex | 0,02 M citrat-Puffer, 1 M sorbitol | RT | 2 h | - | Bösch et al 2016 |
| *U. maydis* | 100 mg/mL Glucanex | 0,02 M citrat-Puffer, 1 M sorbitol | RT | 2 h | - | Bösch et al 2016 |
| *U. maydis/ A. niger* | 20 mg/mL Glucanex<br>0,015 U/mL Chitinase | 0,02 M citrat-Puffer, 1 M sorbitol | RT | up to 20 h | - | Bösch et al 2016,<br>de Bekker et al 2009 |
| *U. maydis/ A. niger* | 20 mg/mL Glucanex<br>0,3 U/mL Chitinase | 0,02 M citrat-Puffer, 1 M sorbitol | RT | 2 h | +/- | Bösch et al 2016,<br>de Bekker et al 2009 |
| *S. indica* | 20 mg/mL Glucanex | 0,02 M MES, 0,05 M CaCl2, 1,33 M sorbitol | RT, 28°C, 37°C | up to 5h | - | Zuccaro et al 2009 |
| *S. indica* | 100 mg/mL Glucanex | 0,02 M MES, 0,05 M CaCl2, 1,33 M sorbitol | RT, 28°C, 37°C | up to 5 h | - | Zuccaro et al 2009 |
| *U. bromivora* | 10 mg/mL Glucanex<br>5 mg/mL Yatalase | 0,02 M MES, 1 M MgSO4 | RT | up to 1,5 h | - | Rabe et al 2016 |
| ***U. bromivora*** | **20 mg/mL Glucanex**<br>**10 mg/mL Yatalase** | **0,02 M MES, 1 M MgSO4** | **RT** | **up to 1,5 h** | **+** | **Rabe et al 2016** |

**Figure S1**

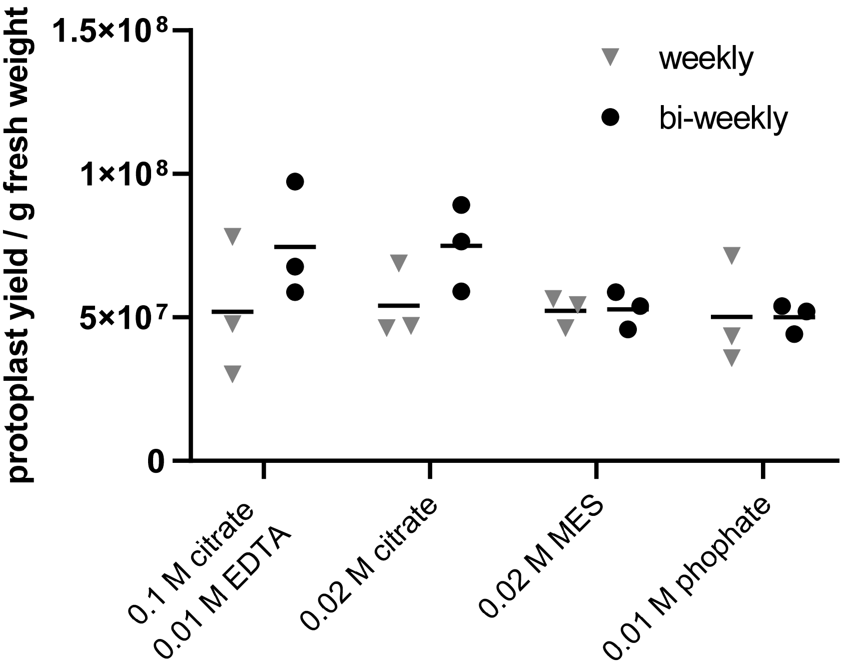


**Fig. S1: Protoplast yields in different buffers are influenced by the sub-culturing scheme.** *T. thlaspeos* LF1 cultures were grown to an OD600<0.8 in an either weekly or bi-weekly splitting rhythm. To test protoplasting efficiency, hyphae were filtered and incubated in different buffers all supplemented with 1.2 M MgSO4 and 10 mg/ml Yatalase + 20 mg/ml Glucanex for 60 mins at RT. There are no significant differences between buffers or splitting rhythm of the culture, but cells sub-cultured bi-weekly yield more protoplasts in citrate buffers.

### Figure S2

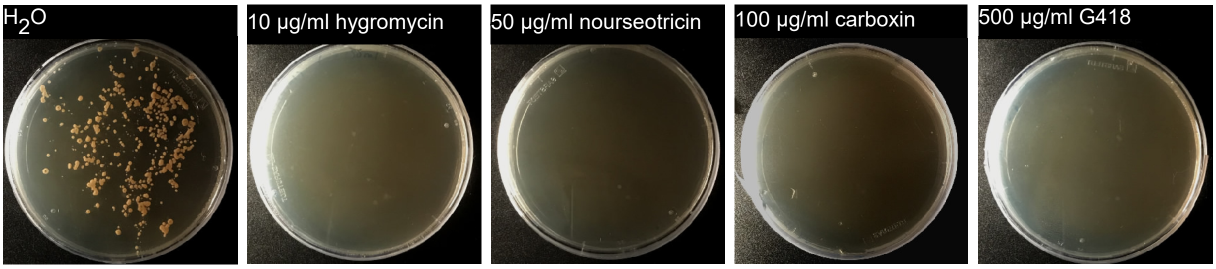


**Figure S2: Sensitivity of *T. thlaspeos* towards different antibiotics.** LF1 was grown on YL 0.6% plant agar plates with and without antibiotics for 16 days. For hygromycin and nourseotricin dilution series were tested and 10 µg hygromycin and 50 µg nourseotricin were identified as minimal inhibitory concentrations to prevent growth. Carboxin inhibits growth at 100 µg/ml and G418 at 500 µg/ml. Both can be used to establish additional resistance markers.

### Figure S3

```
UmSDH2 sen  MSLFNVSNGL RTALRPSVAS S----SRVAA FSTTAAARLA TPTSD-NVGS SGKPQHLKQF KIYRW 60
UmSDH2 res  .......... .......... .----..... .......... .....-.... .......... ..... 60
THTG_04978  ....-AG.T. .A.VQ.V.SG .RSSIA.T.. .....L.Q.. .....S.... .....K..A. ..... 64

UmSDH2 sen  NPDKPSEKPR LQSYTLDLNQ TGPMVLDALI KIKNEIDPTL TFRRSCREGI CGSCAMNIDG VNTLA 125
UmSDH2 res  .......... .......... .......... .......... .......... .......... ..... 125
THTG_04978  .....A.... ..T....... .......... .....V..S. .......... .......... ..... 129

UmSDH2 sen  CLCRIDKQND TKIYPLPHMY IVKDLVPDLT QFYKQYRSIE PFLKSNNTPS EGEHLQSPEE RRRLD 190
UmSDH2 res  .......... .......... .......... .......... .......... .......... ..... 190
THTG_04978  ......REKE S......... V......... .......... .....K.P.A Q......... ..... 194

UmSDH2 sen  GLYECILCAC CSTSCPSYWW NQDEYLGPAV LMQAYRWMAD SRDDFGEERR QKLENTFSLY RCHTI 255
UmSDH2 res  .......... .......... .......... .......... .......... .......... ..L.. 255
THTG_04978  .......... .......... .......... .......... .....T...K T......... A.R.. 259

UmSDH2 sen  MNCSRTCPKN LNPGKAIAQI KKDMAVGAP- KASERPIMAS S 295
UmSDH2 res  .......... .......... .........- .......... . 295
THTG_04978  ...T.....S ...A....T. ..E.ST...A .STD....P. N 300
```

**Fig. S3: Sequence comparison of the succinate dehydrogenase.** *Um*Sdh2 (UMAG_00844) exists in two isoforms that differ in only one amino acid (marked in grey). The carboxin-resistant version carries a leucine instead of a histidine in position 253 (Keon et al., 1991 [41]). *T. thlaspeos* has a homolog that has 82 % amino acid similarity, but has an arginine in the resistance-mediating position. This difference might explain the lower sensitivity against carboxin compared to *U. maydis*.

### Figure S4

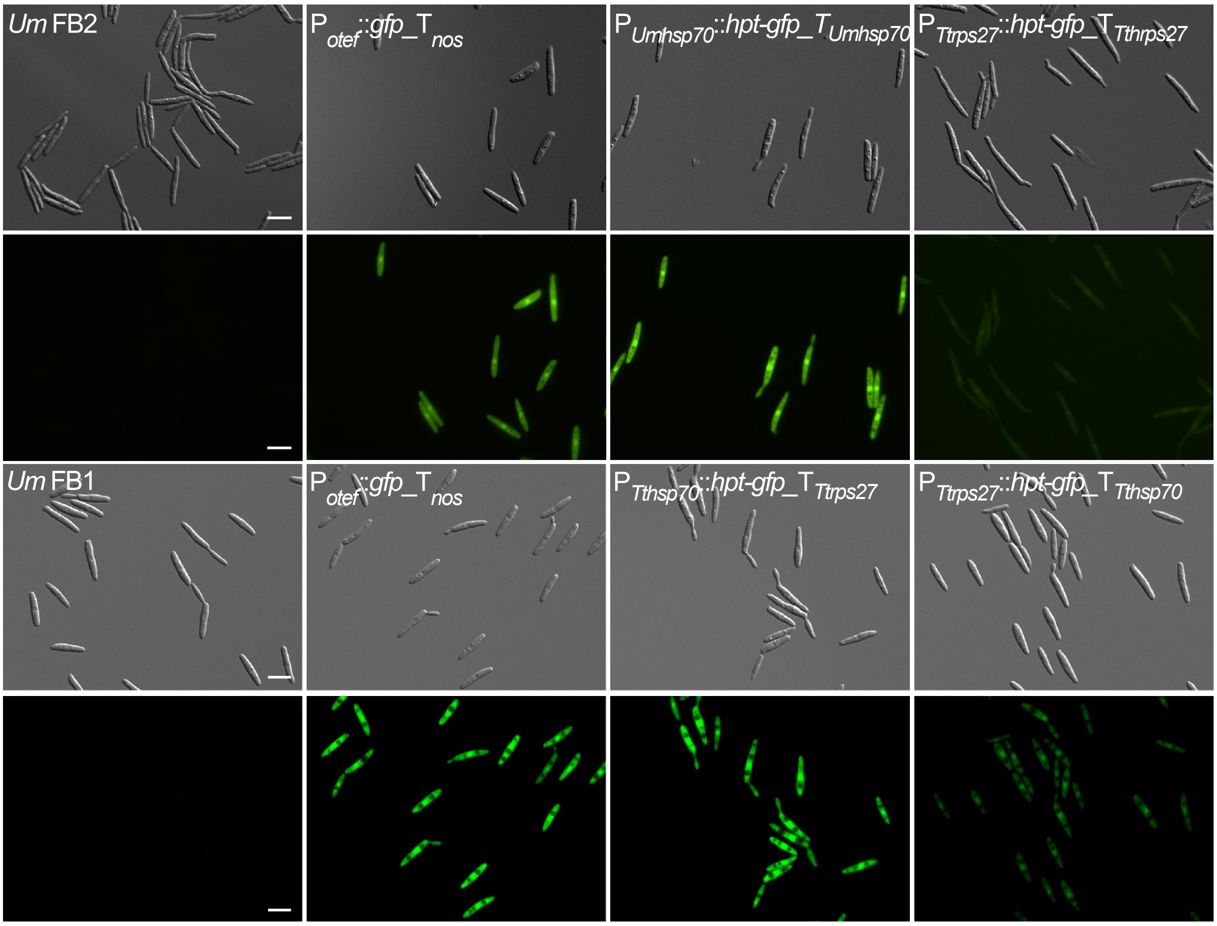


**Figure S4: Verification of resistance-reporter constructs in *U. maydis*.** Reporter constructs containing a fusion of hygromycin resistance (*hpt*) and the fluorescent marker *gfp* under the control of two candidate promoter regions derived from the *T. thlaspeos* genome were tested in *U. maydis* wild-type strains. Upon transformation of the linearized construct, it randomly integrates into the genome. Protein accumulation is visualized by green fluorescence. Gfp-fluorescence in the strains carrying the promoter of *Tthsp70* was stronger compared to the stably integrated construct under the control of a strong, synthetic promoter (P*otef*). Expression under the promoter region of the ribosomal gene *rsp27* was weaker, but the cells still were resistant to hygromycin. This confirms that the fusion protein is active and both promoters can be used, albeit with different strength. Scale bar: 10 µm.

Journal of Fungi

MDPI

**Table S2** Transformation constructs used for the generation of *T. thlaspeos* strains. * indicates that the plasmid was part of the plasmid mix used in the first transformation.

| Vector | Expression cassette | | | | | | Insertion locus | Selective marker | Enzyme |
|---|---|---|---|---|---|---|---|---|---|
| | Promoter | ID | Gene | Terminator | ID | 1 kb flanks | | | |
| pUMa2732* | Um Phsp70 | UMAG_03791 | hpt-gfp | Um Thsp70 | UMAG_03791 | - | ectopic | hygromycin | SspI HF |
| pUMa2790* | Tt Phsp70 | THTG_01007 | hpt-gfp | Tt Thsp70 | THTG_01007 | - | ectopic | hygromycin | SspI HF |
| pUMa2792* | Tt Prps27 | THTG_04331 | hpt-gfp | Tt Trps27 | THTG_04331 | - | ectopic | hygromycin | SspI HF |
| pUMa3030* | Tt Phsp70 | THTG_01007 | hpt-gfp | Tt Trps27 | THTG_04331 | - | ectopic | hygromycin | SspI HF |
| pUMa3031* | Tt Prps27 | THTG_04331 | hpt-gfp | Tt Thsp70 | THTG_01007 | - | ectopic | hygromycin | SspI HF |
| pUMa3576 | Tt Phsp70 | THTG_01007 | hpt-mcherry | Tt Thsp70 | THTG_01007 | - | ectopic | hygromycin | SspI HF |
| pUMa3886 | Tt Phsp70 | THTG_01007 | hpt-gfp | Tt Thsp70 | THTG_01007 | *pra1* (THTG 02790) | *pra1* | hygromycin | SspI HF |

### Protoplasts of *Thecaphora thlaspeos*

- Inoculate 100 ml of YMPG to an $OD_{600}$ of 0,075 with exponentially growing *T. thlaspeos* culture in a 500 ml baffled flask
- Incubate 18°C, shaking at 200 rpm for 3-4 days until $OD_{600}$ reaches 0,6-0,8
- Harvest cells in a cell strainer (40 µm pore size)
- Wash with 20 ml citrate buffer
- Dissolve 20 mg/ml Glucanex and 10 mg/ml Yatalase in citrate buffer and filter-sterilize dissolved enzyme (9 ml enzyme solution /100 ml culture)
- Transfer the filaments into a 50 ml falcon tube and add 9 ml enzyme solution
- Incubate for 30-60 min at RT → formation of protoplasts is visible already after 10 min
- Add 15 ml of citrate buffer and aliquot in 15 ml falcon tubes, 6 ml each
- Overlay with 5 ml trapping buffer (keep everything on ice from now on)
- Spin 15 min 4°C 4863 g in a swing out rotor
- Collect interphases in a 50 ml falcon
- Mix at least with an equal volume of cold STC buffer
- Spin 10 min 4°C 4863 g in a swing out rotor
- Take off supernatant und re-suspend the pellet in 500µl cold STC buffer
- Prepare 100 µl aliquots in 2 ml reaction tubes
- Proceed with transformation immediately

### Transformation of *T. thlaspeos* protoplasts

- Prepare antibiotic bottom-plates (can be prepared the day before):
    - boil up YMPG-REG agar & cool down to 60°C
    - add antibiotic
    - pour 15 ml/plate
    - let it solidify
- Add 15 ml YMPG-REG without antibiotic (prior to transformation)
- Add 1 µl heparin in each transformation tube
- Add linearized DNA (5 µg)
- Incubate 10 min on ice
- Add 500 µl STC/PEG (mix gently)

- Incubate 15 min on ice
- Disperse your transformation on 2 YMPG-REG plates and streak out gently!
- Incubate RT (18°C), wrap with parafilm once they are dry (usually 24 h later)

### Media & solutions

| **YEPSlight** | <u>800 ml</u> |
|---|---|
| 1.0 % (w/v) Yeast-Extract (Difco) | 8.0 g |
| 0.4 % (w/v) Bacto™-Peptone (Difco) | 3.2 g |
| 0.4 % (w/v) sucrose | 3.2 g |

- Dissolve in MilliQ water, aliquot and autoclave for 5 min at 121 ° C

| **1x citrate buffer** | | <u>500 ml</u> |
|---|---|---|
| 0.1 M | trisodium citrate 2x $H_2O$ | 14.75 g |
| 0.01 M | EDTA (0.5M Stock) | 10.00 ml |
| 1.2 M | $MgSO_4$ 7x $H_2O$ | 97.88 g |

- Dissolve citrate and $MgSO_4$ in MilliQ $H_2O$
- Then adjust pH to 5,8 with citric acid solution
- Fill up to 490 ml with MilliQ $H_2O$
- Add EDTA
- Autoclave for 5 min at 121 ° C or filter sterilize

| **1x Citric acid solution** | | <u>200 ml</u> |
|---|---|---|
| 0.1 M | citric acid | 4.20 g |
| 1.2 M | $MgSO_4$ 7x $H_2O$ | 29.58 g |

| **Trapping buffer** | <u>500 ml</u> |
|---|---|
| 0.6M sorbitol | 54.65 g |
| 0.1M Tris/HCl (1M Tris/HCl pH7) | 50.00 ml |

| **YMPG-REG** | <u>800 ml</u> |
|---|---|
| 0.3 % (w/v) Yeast-Extract (Difco) | 2.4 g |
| 0.3 % (w/v) Malzextrakt | 2.4 g |
| 0.5 % (w/v) Bacto™-Pepton (Difco) | 4.0 g |
| 1.0 % (w/v) Glucose | 8.0 g |
| 1.0 M sucrose | 273.8 g |

- In MilliQ water
- Solid medium:
  - + 1.0 % phytagel (4,0 g for 400 ml) or
  - + 0.6 % plant agar (2,4 g for 400 ml
- Aliquot and autoclave for 5 min at 121 °C.
